# Reduced firing rates of pyramidal cells in frontal cortex of APP/PS1 can be restored by acute treatment with levetiracetam

**DOI:** 10.1101/739912

**Authors:** Jan L. Klee, Amanda J Kiliaan, Arto Lipponen, Francesco P. Battaglia

## Abstract

In recent years aberrant neural oscillations in various cortical areas have emerged as a common physiological hallmark across mouse models of amyloid pathology and patients with Alzheimer’s disease. However, much less is known about the underlying effect of amyloid pathology on single cell activity. Here, we used high density silicon probe recordings from frontal cortex area of 9 months old APP/PS1 mice to show that resting state Local Field Potential (LFP) power in the theta and beta band is increased in transgenic animals, while single cell firing rates, specifically of putative pyramidal cells, are significantly reduced. At the same time, these sparsely firing pyramidal cells phase-lock their spiking activity more strongly to the ongoing theta and beta rhythms. Furthermore, we demonstrated that the anti-epileptic drug, levetiracetam, can restore principal cell firing rates back to control levels. Overall, our results highlight reduced firing rates of cortical pyramidal cells as a symptom of amyloid pathology and indicate that lifting cortical inhibition might contribute to the beneficial effects of levetiracetam on AD patients.

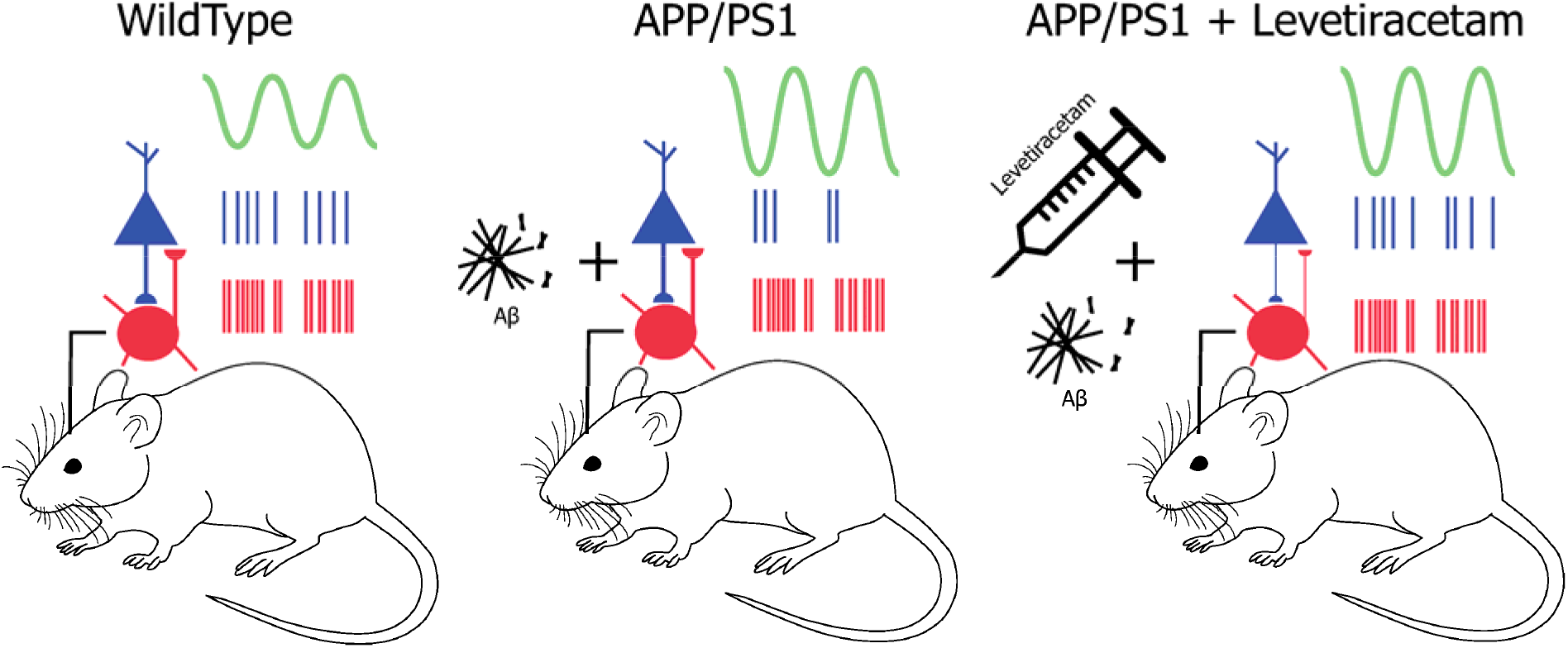

## 1. Introduction

Network hyper-synchrony and altered neural oscillation have been suggested to contribute to the pathophysiology of Alzheimer’s disease (AD). A better understanding of altered neural oscillations in AD could therefore provide a target for pharmacological interventions. Even though amyloid plaques and related neuronal loss are among the most significant findings in the post mortem brain of AD patients, the amount of plaques does not correlate with the severity of dementia (Nagy et al., 1995), and the removal of plaques does not lead to an improvement of memory (Holmes et al., 2008). Intriguingly, amyloid accumulation seems to cause excitatory-inhibitory imbalance at the synaptic level, triggering abnormal patterns of single cell, as well as neuronal network activity and epileptiform discharges (Minkeviciene et al., 2009). Simultaneously with amyloid plaque formation, both hyper- and hypoactive neurons emerge in the hippocampus but also in the cortical areas (Busche et al., 2008; Heggland et al., 2019; Xu et al., 2015). Less is known about which subpopulations of cells are affected by amyloid accumulation but it has been proposed that this pathological process is related to persistently decreased resting membrane potential in neocortical pyramidal neurons (Minkeviciene et al., 2009). Similarly, it has been reported that basic biophysical properties of pyramidal neurons in frontal cortex are intact but external stimulation of these neurons revealed hyper-excitability, indicating a combination of both intrinsic electrical and synaptic dysfunctions as mechanisms for activity changes (Kellner et al., 2014). Whether these findings apply to in vivo unanesthetized mice, needs to be verified.

On the level of neuronal assemblies, the most prominent finding related to amyloid pathology is abnormally high LFP (Local Field Potential) power over a broad frequency range during wide variety of behavioral states (Goutagny et al., 2013; Gurevicius et al., 2013; Jin et al., 2018; Pena-Ortega et al., 2012; Verret et al., 2012), which can lead to epileptiform synchronous discharges and generalized seizure activity (Gurevicius et al., 2013; Jin et al., 2018; Lam et al., 2017; Minkeviciene et al., 2009; Palop and Mucke, 2016; Vossel et al., 2013). The mechanism by which aberrant single cell activity changes into generalized epileptiform activity of neuronal ensembles is not clear. Traditionally, epileptic seizures have been characterized as hypersynchrony of large neuronal populations leading to the epileptiform state (Jiruska et al., 2013). However, this view has been challenged by the finding that, during epileptic seizures, there are both increases and decreases in firing rates of neurons and many units remaining unchanged (Schevon et al., 2012; Truccolo et al., 2011; Wyler et al., 1982). Furthermore, single cell activity outside the periods of seizures and areas of epileptic foci is more heterogeneous and unsynchronized, and not well characterized (Keller et al., 2010; Truccolo et al., 2011). In addition, to our knowledge it is not known if amyloid accumulation affecting EEG power in broad frequency band is also entraining single cell activity.

In recent years, antiepileptic pharmacological treatments to balance altered neuronal activity as a consequence of amyloid accumulation have become of interest (Cumbo and Ligori, 2010; Ziyatdinova et al., 2015, 2011). Acute dose of levetiracetam in a animal model reduces abnormal EEG spiking activity in the cortex and in the hippocampus for 18 hours after administration (Sanchez et al., 2012). Notably, sub-chronic treatment with levetiracetam has been shown to boost abnormal hypoactivation in the entorhinal cortex in people with amnestic mild cognitive impairment (aMCI) while simultaneously improving working memory performance (Bakker et al., 2012). Currently, this treatment is under clinical testing although the basic mechanisms of action of levetiracetam in AD patients is not well understood (Bakker et al., 2015). In animal models overexpressing amyloid protein, chronic levetiracetam treatment suppresses the amyloid accumulation and improves hippocampal dependent spatial memory (Sanchez et al., 2012). However, levetiracetam has multiple plausible molecular targets including voltage-gated ion currents, synaptic vesicle proteins and the glutaminergic system (Surges et al., 2008), and it is not known which ones are relevant for alleviating AD symptoms.

To investigate these questions, we recorded both LFP and single cell activity in head-fixed APP/PS1 mice and analyzed the effect of acute levetiracetam treatment. Surprisingly, we found that while LFP oscillations showed a power increase in the theta and beta frequency range as previously reported, frontal cortex pyramidal cell firing rates were significantly reduced in APP/PS1 mice. At the same time, the sparsely firing pyramidal cells phase-locked more strongly to the ongoing theta rhythm, revealing a non-trivial relationship between increased oscillations and micro-circuit effects. Levetiracetam specifically elevated pyramidal cell firing rates in APP/PS1 mice back to control levels and decoupled pyramidal cells and interneurons as shown by decreased pairwise correlations.

In combination, our results indicate that reduced firing rates of cortical pyramidal cells emerge as a symptom of amyloid pathology and that reducing inhibition might be a viable approach to improve brain function in AD.

## 2. Material and Methods

### 2.1 Animals

For the present study we used male APP_swe_/PS1_dE9_ (APP/PS1) transgenic mice and their age-matched wild type (wt) littermates. The APP/PS1 founder mice were originally obtained from John Hopkins University, Baltimore, MD, USA (D. Borchelt and J. Jankowsky, Department of Pathology) and a colony was first established at the University of Kuopio, Finland and thereafter a colony was bred at the Central Animal Facility at Radboud university medical center, The Netherlands. The mice were created by co-injection of chimeric mouse/human AβPPswe (mouse AβPP695 harbouring a human Aβ domain and mutations K595N and M596L linked to Swedish familial AD pedigrees) and human PS1-dE9 (deletion of exon 9) vectors controlled by independent mouse prion protein promoter elements. The two transfected genes co-integrate and co-segregate as a single locus (Jankowsky et al., 2001). This line was originally maintained on a hybrid background by backcrossing to C3HeJ×C57BL/6J F1 mice (so-called pseudo F2 stage). For the present work, the breeder mice were backcrossed to C57BL/6J for 15 generations to obtain mice for the current study. The animals were 9 months old at the start of the experiment. All animals were group housed until the first surgery after which they were individually housed to prevent damage to the implants. Throughout the experiment the animals received food and water ad libitum and were maintained on a 12 hours light/dark. Recordings were performed during the light period. All experiments were approved by the Dutch governmental Central Commissie Dierproeven (CCD) (10300) and conducted in accordance with the ARRIVE guidelines (Kilkenny et al., 2012).

### 2.2 Surgical preparation for head-fixed electrophysiological recordings

Animals were anesthetized using isoflurane (0.8 – 1 l/min, 1.5-2 %). and placed in a stereotaxic frame. At the onset of anesthesia, all mice received subcutaneous injections of carprofen (5 mg/kg) as well as a subcutaneous lidocaine injection through the scalp. The animals’ temperature was maintained stable for the duration of the surgical procedure using a heating blanket. The level of anesthesia was checked during operation by pedal reflex. We exposed the skull and inserted a skull screw over the cerebellum to serve as combined reference and ground. We then placed a custom made, circular head-plate for head-fixation evenly on the skull and fixated it with dental cement (Super-Bond C&B, Sun Medical, Shiga, Japan). A small craniotomy was drilled over left frontal cortex at +1.78mm anterior and 0.4 mm lateral to bregma and the exposed skull was covered with a silicone elastomer until the first recording. All mice were given at least 2 days to recover from the surgery.

### 2.3 Electrophysiological recordings in frontal cortex of head-fixed mice

The head-fixation setup consisted of two rods that were screwed onto either side of the implanted head-plate and fixated the mice in place on top of a Styrofoam ball that functioned as an air-supported spherical treadmill and allowed us to read out the movement of the animals in a subset of recording sessions. All animals were slowly habituated to head-fixation by placing them in the setup for 3 sessions of 10 minutes for two days prior to the first recording day.

Two hours prior to each recording session, we injected the animals with either 200 mg/kg i.p. levetiracetam, (Sigma-Aldrich, Taufkirchen, Germany), a dose shown to reduce abnormal hyperactivity EEG effectively, or saline (Doheny et al., 1999; Sanchez et al., 2012). We then alternated these injections for 4 consecutive days to be able to record two sessions per animal and condition (Doheny et al., 1999). The experimenter was blind to both the genotype as well as to the type of injection.

At the start of each recording session, we placed the mice in head-fixation and removed the silicon elastomer cover to expose the craniotomy over the frontal cortex. We then used a micromanipulator (Thorlabs, Newton, New Jersey, USA) to acutely insert a 128 channel silicon (IMEC, Leuven, Belgium) probe −1.5mm and −2.5mm deep into the frontal cortex (Dimitriadis et al., 2018). At each depth, we waited about 15 minutes for the tissue to settle. We then performed 10 minutes of broadband recordings.

Electrophysiological signals were filtered between 1 and 6000 Hz, digitized at 30 kHz using 2 digitizing headstages (RHD2164 Amplifier Board, Intan Technologies, Los Angeles, California, USA) and acquired with an open-ephys recording system. After each recording session, we retracted the silicon probe and placed a new silicon cover on the skull before releasing the animals back to their respective home cages.

### 2.4 Histology

At the end of the experiment, the animals were deeply anesthetized with pentobarbital (Nembutal, pentobarbitalsodium 60 mg/ml, 65 mg/kg, i.p.) and perfused with 0.1 M phosphate buffer (pH = 7.4.) followed by 4% paraformaldehyde solution for 9 min at 10 mL/min. The brain was removed and left for immersion postfixation for 4 h in 4% paraformaldehyde (pH = 7.4) at 4°C. The brains were thereafter stored 0.1. phosphate buffer (pH = 7.4) at 4°C until slicing. Coronal sections (thickness 35 μm) were cut with a freezing slide microtome. Every second section containing track from the recording electrode were stained for the N-terminal human Aβ-specific antibody W02 (Genetics, Switzerland) to visualize diffuse amyloid deposits as described previously (Hooijmans et al., 2007) to verify the amyloid pathology in the transgenic animals (Fig. 1.). Additionally, we performed cresyl-violet staining to verify the anterior-posterior coordinates of the recording sites.

**Figure 1.**
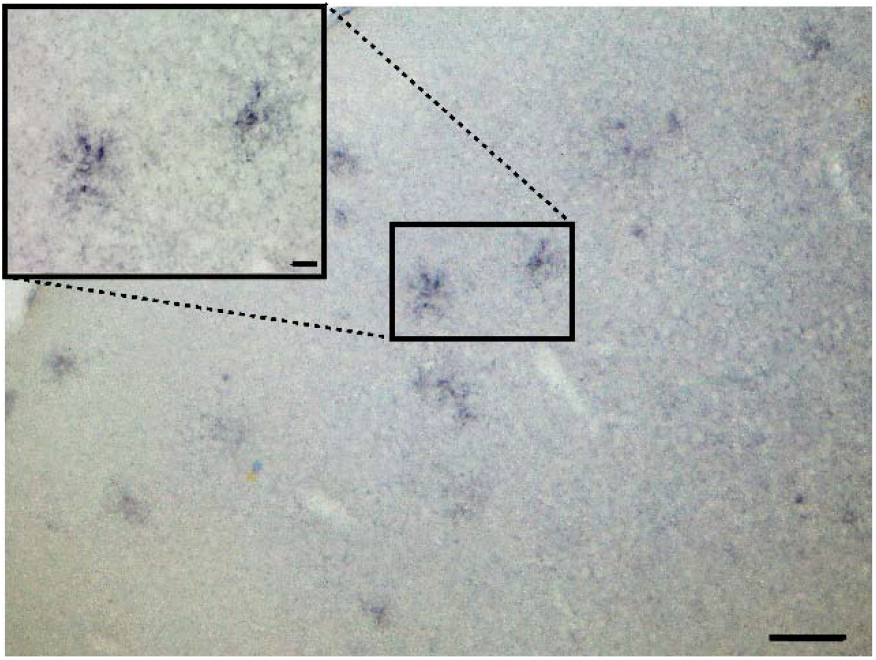
A representative histological section from a transgenic APP/PS1 mouse. W02 antibody staining for Aβ deposits revealed plaques in the frontal cortex. Scale bars 100 μm in the main figure and 20 μm in the zoom figure.

### 2.5 Data analysis

For LFP analysis, we only used the central channel of our recording electrode. We down-sampled the broadband signal to 2000 Hz and then low-pass filtered below 500 Hz. Time-frequency analysis was performed via Morlet wavelet convolution based on the fast Fourier transform (Cohen, 2014). To minimize the effect of behavioral changes on our results, we split each recording session into 4-s bins and focused our subsequent analysis on then 20 bins with the highest theta power. Significant differences in the power spectra were calculated using a permutation test at each frequency.

To identify single units, data were automatically spike sorted with Kilosort (Pachitariu et al., 2016) (https://github.com/cortex-lab/Kilosort) and then manually inspected with the phy software (https://github.com/kwikteam/phy). All following analyses were performed using custom written Matlab scripts. As a post-hoc criterion for the quality of our spike sorting, we computed the auto-correlograms of each putative single unit and only further analyzed those with less than 2% of spikes within the physiological refractory period of 2ms of each other.

For each unit we then computed the peak-to-peak amplitude of the action potential waveforms as well as the width at 30% of the negative peak. We fed these values into Gaussian-mixture model to classify single units into putative interneurons with narrower spike waveforms and putative pyramidal cells with broader spike waveform (see Fig. 3 for cut off criteria) (Stark et al., 2013). For each unit, we computed the mean firing rate over the 10-min recording sessions and performed a 3-factorial ANOVA with neuron type, genotype and drug condition as independent variables, followed by post-hoc t-tests wherever justified by significant main or interaction effects.

Additionally, we computed the spike-time cross-correlations between all simultaneously recorded neurons in our experiment and performed the same statistical analysis as above (Moore et al., 1966; Perkel et al., 1967a, 1967b). For theta and beta phase locking analysis, we first filtered the raw signal between 4-12 Hz and 12-30 Hz respectively using a zero-lag band-pass filter (filtfilt function in the Matlab software), and then extracted the phase angles using the Hilbert transform. For each putative single cell, we extracted the theta and beta phase angles at each spike and computed the mean resultant length. Significant phase locking was computed using the Rayleigh test for circular uniformity.

## 3. Results

### 3.1 Increased LFP power in 6-26 Hz band in APP/PS1 mice

Studies have consistently reported increased cortical LFP power in theta (4-12 Hz) and beta (13-30 Hz) frequency bands in cortex of APP/PS1 mice at various developmental stages from 4 to 16 months of age (Gurevicius et al., 2013; Jin et al., 2018). However, the relationship between the local field potential and changes in the underlying single cell activity caused by amyloid pathology have not been explored. In this context, we recorded both local field potential as well as single cell activity with a 128-channel silicon probe in frontal cortex of 9-month-old awake, head-fixed APP/PS1 mice, showing amyloid accumulation (Fig. 1) and wildtype littermate controls. In line with our previous results, we found increased theta and beta LFP power between 6 and 26 Hz in the frontal cortex neurons of APP/PS1 mice (APP/PS1 n=4 mice, 6 sessions; WT n=3 mice, 4 sessions; permutation test at each frequency *p*<0.05) (Fig. 2.).

**Figure 2.**
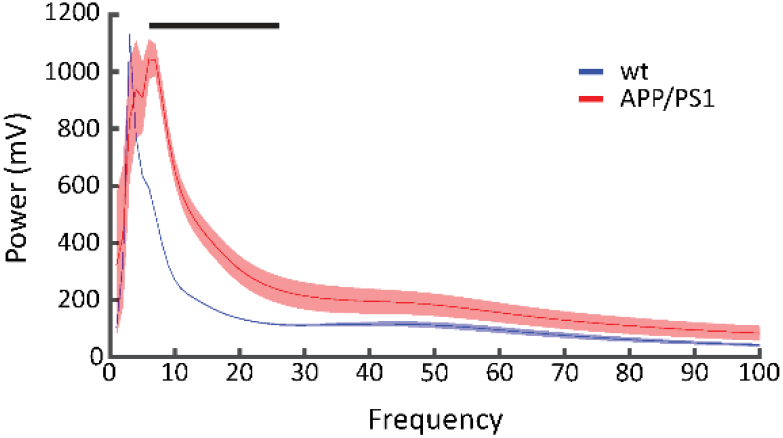
Increased LFP Power in APP/PS1 mice; Population average power from APP/PS1 mice and wildtype controls (APP/PS1 n=4 mice, 6 sessions; WT n=3 mice, 4 Sessions). Black bar indicates significant power differences from 6-28 Hz (permutation test at each frequency p<0.05). Shaded areas indicate SEM.

### 3.2 Reduced single cell firing rates in frontal cortex of APP/PS1 mice

Increased LFP power in combination with consistent reports of epileptiform activity point to network hyperexcitability in APP/PS1 mice but evidence from single cell recordings in these animals is sparse (Gurevicius et al., 2013; Jin et al., 2018). Therefore, we recorded a total of 1705 individual neurons from the frontal cortex of APP/PS1 mice under various conditions. Surprisingly, we found that APP/PS1 mice showed overall reduced firing rates compared to the wildtype controls (*t*-test, p = 0.011).

To further investigate whether this effect was specific to a certain neuron type, we classified the recorded single cells as either putative pyramidal cells or interneurons according to their action potential waveform (Stark et al., 2013). This analysis revealed that the amyloid pathology related reduction of firing rates in APP/PS1 mice was specific to putative pyramidal cells (ANOVA genotype-neurontype interaction F(1,1427)=5.99, p=0.01; post hoc *t*-test for APP/PS1 vs WT pyramidal cells, p=0.006, ^~^22% reduction in mean firing rates) while interneurons were not significantly affected (post hoc *t*-test for APP/PS1 vs WT interneurons, p=.97 ^~^.5% increase in mean firing rates (Fig. 3C. and Fig. 4A.).

**Figure 3.**
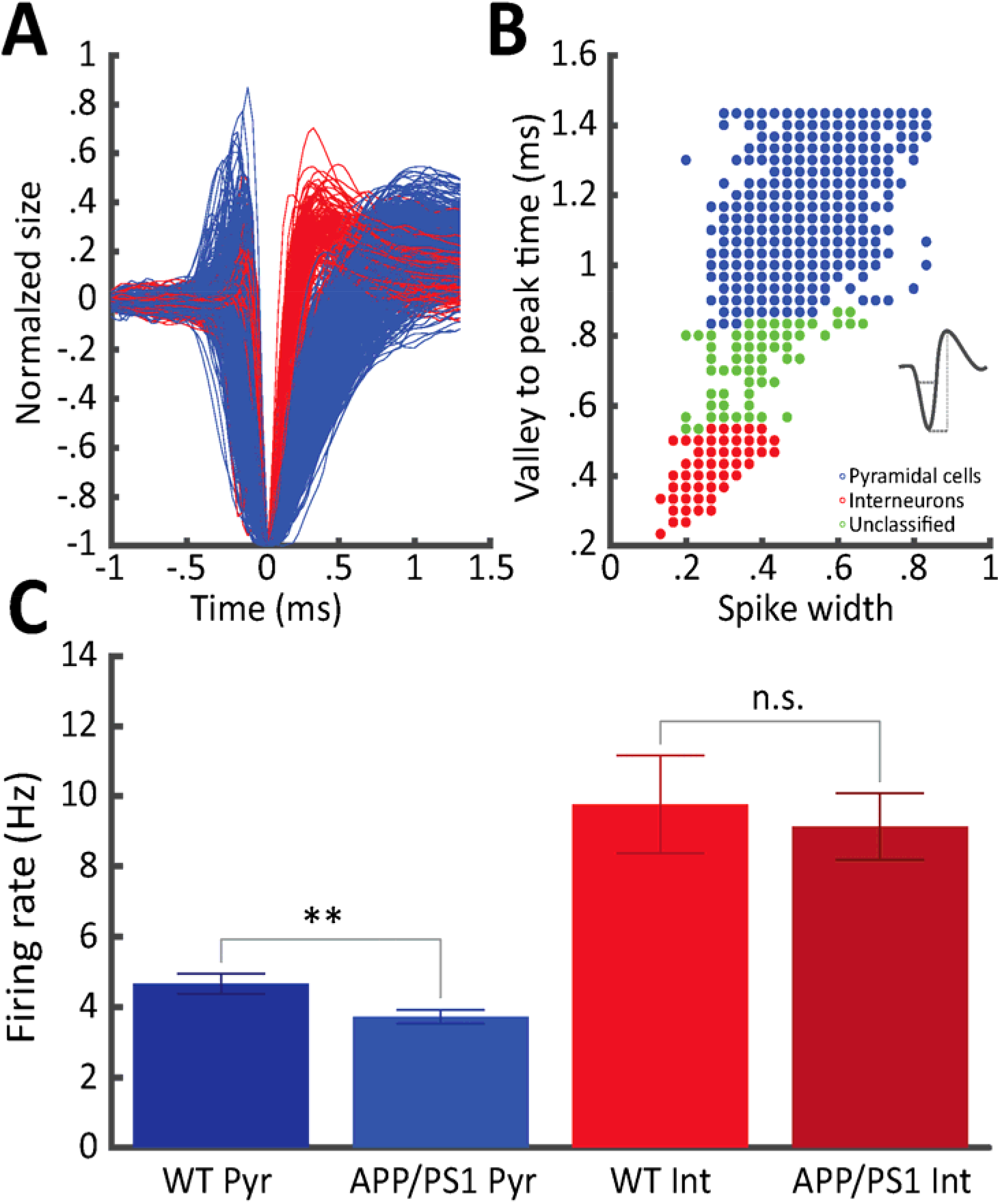
Decreased pyramidal cell firing rates in APP/PS1 mice A) Average action potential waveforms of all recorded putative pyramidal cells (blue) and interneurons (red) B) Spike width to valley to peak ratio of all recorded single cells color-coded according to Gaussian mixture model classification. C) Average firing rates of putative pyramidal cells and interneurons in wild-type controls and APP/PS1 mice (n=228 (WT/Pyr); 386 (APP/PS1/Pyr); 35 (WT/Int); 45 (APP/PS1/Int); ** indicates p<0.01). Error bars represent SEM.

**Figure 4.**
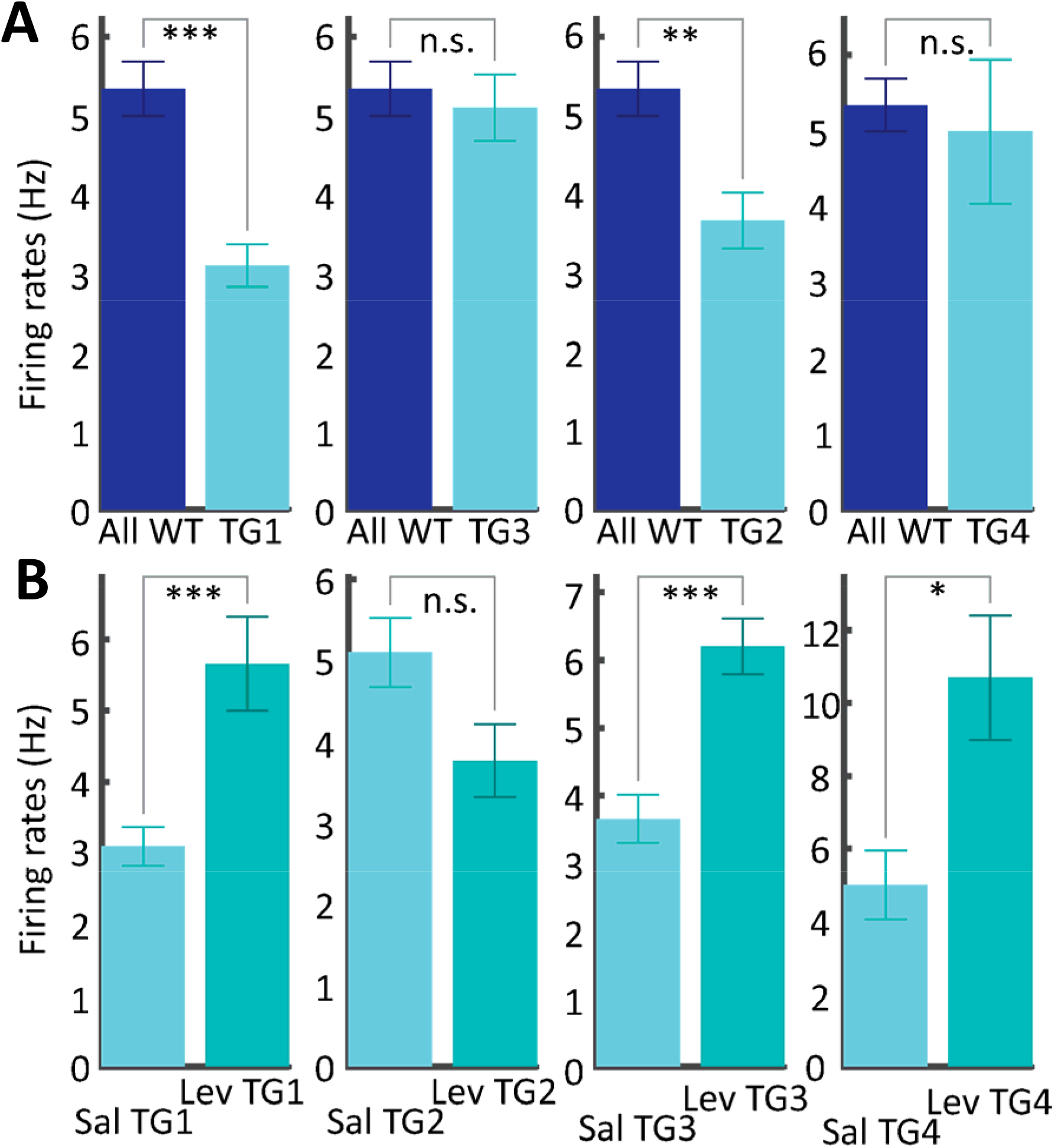
Firing rate differences in individual animals. A) Average pyramidal cell firing rates are consistently lower in APP/PS1 mice compared to wildtype controls. B) Levetiracetam increases pyramidal cell firing rates in APP/PS1 mice compared to wildtype controls.

### 3.3 Increased theta and beta phase locking of pyramidal cells in APP/PS1 mice

Our analysis of LFP and single cell activity in APP/PS1 mice points towards non-trivial relationship between increased local field potential oscillations in the theta and beta range and the underlying single cell firing rates. To shed light on this issue, we made use of our concomitant LFP and unit recordings and analyzed the entrainment of putative pyramidal cells to the predominant ongoing LFP oscillation in the 4-12 Hz theta and beta range (Fig. 2. and Fig. 5.). To this end, we filtered the raw LFP signal between 4-12 Hz and 12-30 Hz, respectively, and then computed the phase angles using the Hilbert transform. For each putative single cell, we extracted the theta and beta phase angles for each spike and computed the mean resultant length. We found that the overall more sparsely firing pyramidal cells in APP/PS1 mice were significantly more phase-locked to the ongoing theta and beta oscillation than in wild type controls (theta: Wilcoxon Rank sum test, *p*<0.001 (Fig. 5B.); beta: Wilcoxon Rank sum test, *p*<0.001 (Fig. 6A.); n=228 (WT/Pyr); 386 (APP/PS1/Pyr)) Phase locking in the gamma range between 30-100 Hz was not significantly affected (Fig. 6B.).

**Figure 6.**
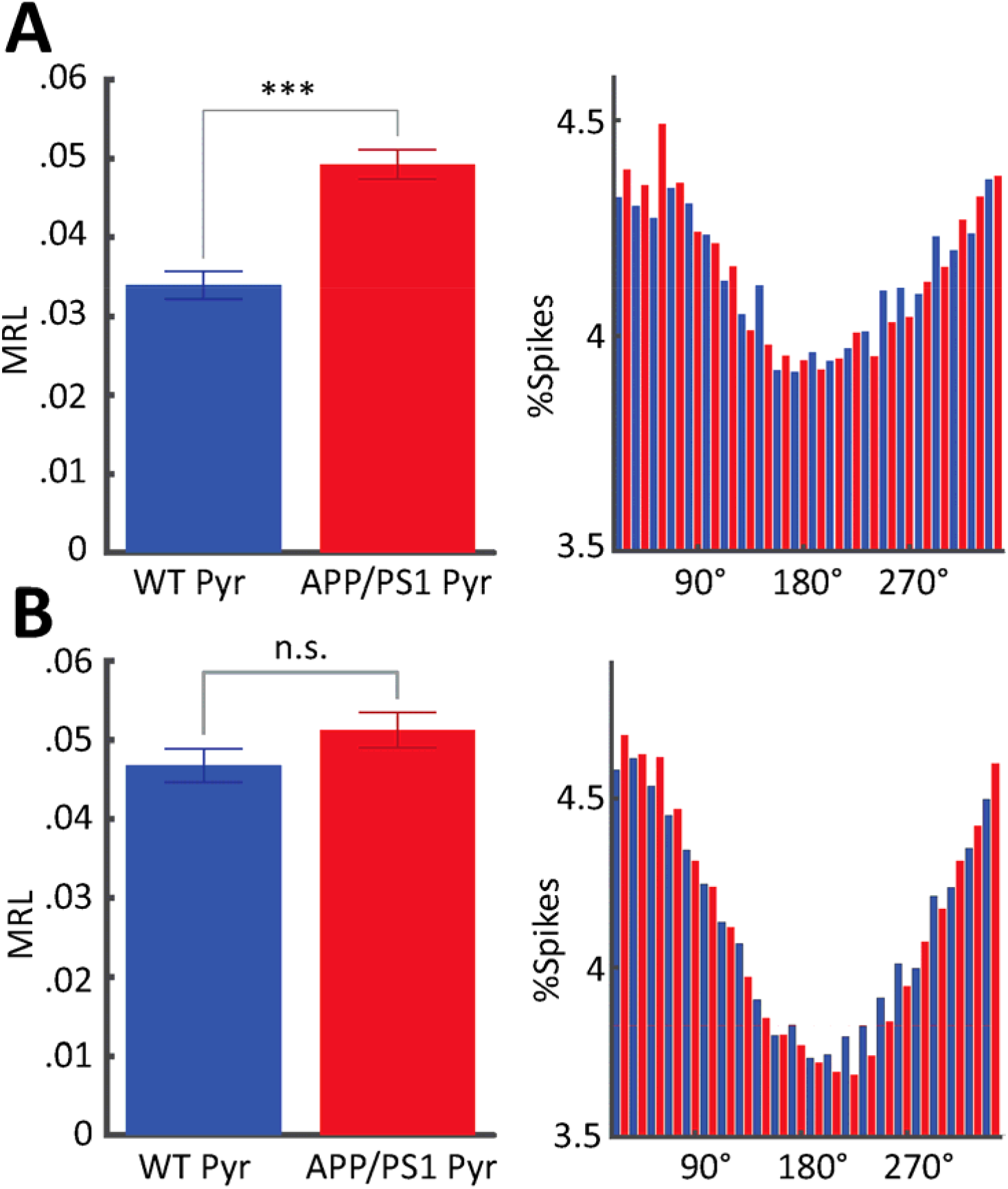
A) Left, APP/PS1 pyramidal cells are significantly more locked to the ongoing beta oscillations than pyramidal cells in wild-type controls (MRL= mean resultant length, ***indicates p<.001, n=228 (WT/Pyr); 386 (APP/PS1/Pyr)). Error bars represent SEM. Right: Distribution of average firing phase of all significantly modulated WT(blue) and APP/PS1 (red) pyramidal cells (Rayleigh test for circular uniformity) B) Left: APP/PS1 pyramidal cells are not significantly more locked to the ongoing gamma oscillations than pyramidal cells in wild-type controls (30-100 Hz). Right: Distribution of average firing phase of all significantly modulated pyramidal cells.

### 3.4 Levetiracetam restore reduced firing rates of single neurons in frontal cortex of APP/PS1 mice

Interestingly, we found that levetiracetam specifically elevated firing rates in frontal cortex neurons in APP/PS1 mice (Fig. 7. and Fig. 4B.) while firing rates in wildtype animals were not significantly affected (genotype by drug interaction F(1,1427)=7.79, *p*=0.005; post hoc *t*-test for APP/PS1 cells + lev vs APP/PS1 cells + sal, *p*<0.001). Importantly, classifying cells into putative pyramidal cells and interneurons according to their action potential waveforms revealed that this effect was specific for pyramidal cells (neuron type by drug interaction F(1,1427)=3.91, *p*=0.048; post hoc *t*-test for APP/PS1 pyramidal cells + lev vs APP/PS1 pyramidal cells + sal, *p*=0.001). Levetiracetam therefore specifically reversed the perturbing effect of amyloid pathology on pyramidal cells in APP/PS1 mice.

### 3.5 Levetiracetam uncouples pyramidal cells and interneurons in APP/PS1 mice

Levetiracetam has been shown to suppress vesicle release (De Smedt et al., 2007a; Surges et al., 2008; Vogl et al., 2012). Increased firing rates of pyramidal cells in APP/PS1 after treatment with levetiracetam therefore seems more likely to be the result of reduced inhibition rather than a direct stimulating effect on pyramidal cells. To find support for this hypothesis, we analyzed the pair-wise correlations of all simultaneously recorded single cells in APP/PS1 mice and focused our attention on pyramidal cell - interneuron correlations (Fig 7). In line with its overall positive effect on single cell firing rates, levetiracetam significantly increased pair-wise pyramidal cell correlations in APP/PS1 mice. However, pyramidal-interneuron correlations were significantly reduced, effectively uncoupling interneurons and pyramidal cells in APP/PS1 mice (three-way interaction between genotype, neuron type and drug F(3,50291)=.643, p=0.002; post hoc *t*-test for APP/PS1 pyramidal cells + lev vs APP/PS1 pyramidal cells + Sal, *p*<0.001; APP/PS1 interneurons+ lev vs APP/PS1 interneurons + sal *p*< 0.001).

**Figure 7.**
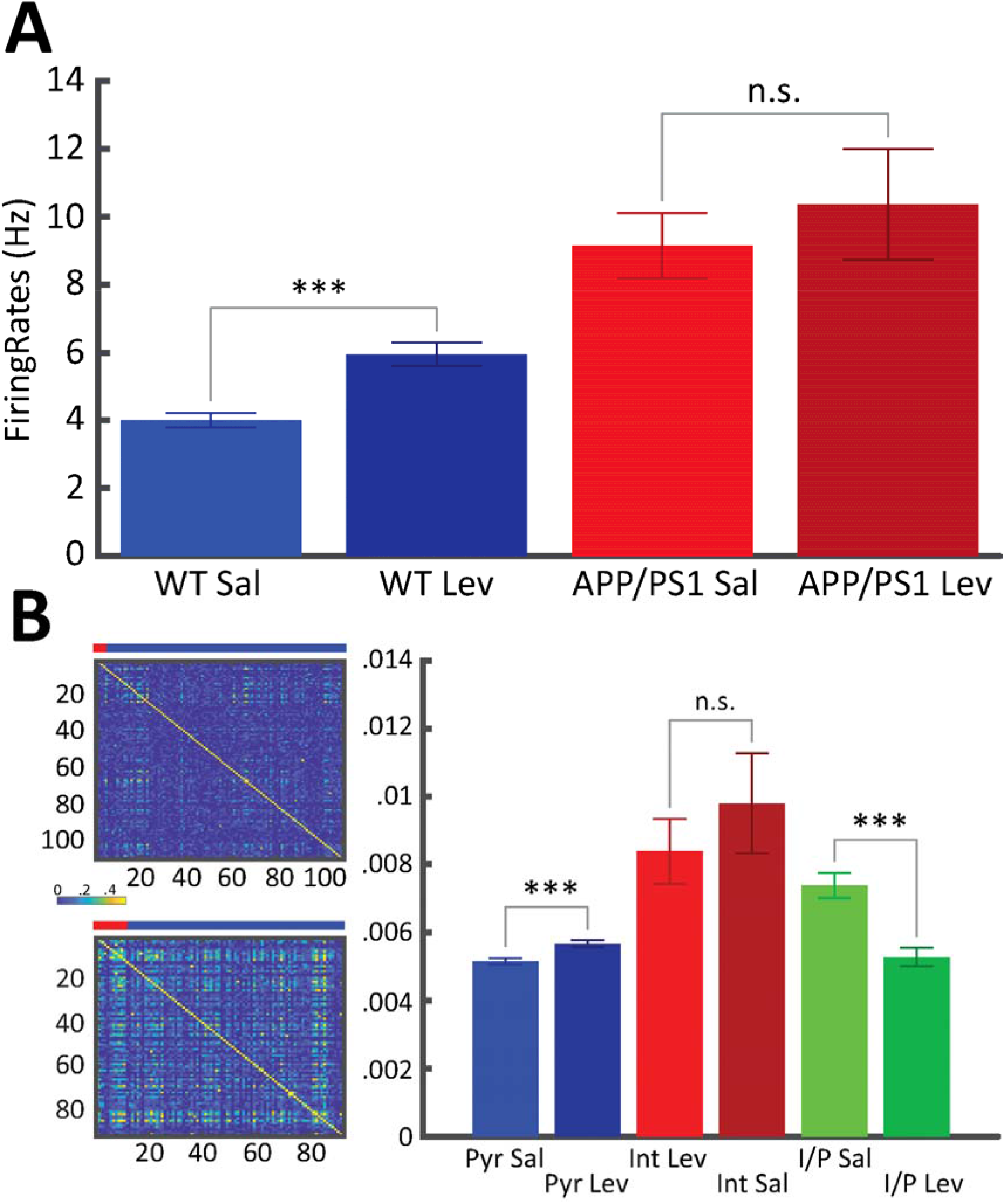
Levetiracetam restores pyramidal cell firing rates in APP/PS1 mice. A) Average firing rates of putative pyramidal cells and interneurons in APP/PS1 mice treated either with saline or levetiracetam injections (n=386 (APP/PS1/Pyr/Sal); 326 (APP/PS1/Pyr/Lev); 45 (APP/PS1/Int/Sal); 39 (APP/PS1/Int/Lev; *** indicates p<0.001). Error bars represent SEM. B) Example sessions of pairwise correlations of simultaneously recorded neurons from one animal with saline (top) and levetiracetam (bottom) injections. Red bar indicates interneurons and blue pyramidal cells. Right, average pairwise correlation of all simultaneously recorded neurons according to cell type.

### 3.5 Levetiracetam uncouples pyramidal cells and interneurons in APP/PS1 mice

Levetiracetam has been shown to suppress vesicle release(De Smedt et al., 2007a; Surges et al., 2008; Vogl et al., 2012). Increased firing rates of pyramidal cells in APP/PS1 after treatment with levetiracetam therefore seems more likely to be the result of reduced inhibition rather than a direct stimulating effect on pyramidal cells. To find support for this hypothesis, we analyzed the pair-wise correlations of all simultaneously recorded single cells in APP/PS1 mice and focused our attention on pyramidal cell - interneuron correlations (Fig 7). In line with its overall positive effect on single cell firing rates, levetiracetam significantly increased pair-wise pyramidal cell correlations in APP/PS1 mice. However, pyramidal-interneuron correlations were significantly reduced, effectively uncoupling interneurons and pyramidal cells in APP/PS1 mice (three-way interaction between genotype, neuron type and drug F(3,50291)=.643, p=0.002; post hoc *t-* test for APP/PS1 pyramidal cells + lev vs APP/PS1 pyramidal cells + Sal, *p*<0.001; APP/PS1 interneurons+ lev vs APP/PS1 interneurons + sal *p*< 0.001).

## 4. Discussion

In line with previous studies, we found that LFP power was increased in 9-month-old APP/PS1 mice in the theta and beta frequency bands between 6 and 26 Hz (Gurevicius et al., 2013; Jin et al., 2018). However, contrary to the prevailing interpretation that increased LFP power indicates hyperactivity on the single cell level in AD mouse models, we found that firing rates, specifically of frontal cortical pyramidal cells, were significantly reduced in APP/PS1 mice compared to wildtype controls (Jiruska et al., 2013). One explanation for these counterintuitive results was our finding that phase-locking of pyramidal cells to the dominant cortical theta and beta oscillations increases and that this can lead to an increase in LFP power. Interestingly, it has been shown previously that GABA levels in frontal cortex of AD mice can rise continuously and peak around 9 months of age, the same age as our cohort (Roy et al., 2018). Therefore, one plausible interpretation of our results is that increased cortical inhibition in APP/PS1 mice leads to a narrower window of opportunity for pyramidal cell spiking. While this would need to be verified in our model, pathologically increased cortical inhibition could explain overall decreased firing rates as well as the more precise alignment of spikes with the ongoing theta and beta oscillations.

Acute systemic injection of levetiracetam reversed the effects of AD pathophysiology and specifically restored pyramidal cell firing rates back to control levels. Levetiracetam is commonly used to treat neuronal hyperactivity during epileptic seizures (Cumbo and Ligori, 2010; De Smedt et al., 2007b). Therefore, it seems unlikely to us that levetiracetam directly acts in an excitatory way on pyramidal neurons. Furthermore, it has been shown previously that levetiracetam reduces vesicle release from presynaptic terminals (Vogl et al., 2012). Therefore, we assume that levetiracetam reduces the release of inhibitory transmitters from cortical interneurons and thereby releases pyramidal cells from inhibition. Along this line of interpretation reduced cortical inhibition could be explained without a change in firing rates of cortical interneurons. The reduced interneuron/pyramidal cell correlation that we demonstrate in Figure 7 is compatible with this view.

Overall, our results point towards reduced excitability of pyramidal cells in APP/PS1 mice which results in reduced firing rates and more phase-locking to the ongoing theta and beta oscillations. Levetiracetam seems to specifically counteract this effect, as it uncouples pyramidal cells from putative inhibitory interneurons and elevates pyramidal cell firing rates. Targeting inhibitory transmission in AD therefore seems to us as a viable new route to potentially reverse network dysfunction in AD.

## Acknowledgements

We thank Prof. Heikki Tanila for helpful comments on the manuscript and Jos Dederen and Vivienne Verweij for excellent technical support. The study was supported by the Säätiöiden post doc - pooli (The Finnish Cultural Foundation) to Dr. Arto Lipponen. Silicon probes were manufactured by IMEC (Leuven, Belgium) with funding from the European Union’s Seventh Framework Programme (FP7/2007-2013) under grant agreement nr. 600925 (NeuroSeeker).

